# The interplay of cortical magnification and perceptual load in the visual processing of task-irrelevant biological motion across the visual field

**DOI:** 10.1101/2023.08.02.551722

**Authors:** Murat Batu Tunca, Ada Dilek Rezaki, Hilal Nizamoğlu, Burcu A. Urgen

## Abstract

Perceptual load theory argues that attention is a limited resource and stimuli cannot be processed if there is insufficient perceptual capacity available. Although attention is known to modulate biological motion processing, whether this modulation differs among different perceptual loads remains unknown. To answer this question, three experiments are conducted in which biological motion is utilized as a task-irrelevant distractor. The first experiment showed that biological motion is processed differently than non-biological motion across different perceptual load conditions. The second experiment investigated the effect of attention on biological motion processing, revealing that higher eccentricities enhance biological motion processing but only when the perceptual load is low. The last experiment investigated the same question but with cortically magnified stimuli. It found that when the stimuli are cortically magnified, the enhancement effect of eccentricity is present regardless of the perceptual load. Overall, the results suggest that perceptual load modulates the processing of task-irrelevant biological motion and interacts with other factors (such as eccentricity) that modulate this processing.

## 1. Introduction

The ability to attend to and further process the selected stimulus is called selective attention. Previous research reveals many factors that affect selective attention, one of which is task demands (Johnston & Dark, 1986). The perceptual load theory of attention is a widely accepted model that explains how attention is highly task-dependent (Lavie, 1995). According to the theory, attention is not an unlimited resource. All humans have a capacity, and each stimulus will fill this capacity partially. The stimuli will compete for our attention, and selective attention plays a top-down role to determine which stimulus will be the primary focus of attention. If the attended stimulus requires a low perceptual capacity to be processed, there will be spared capacity which will be filled with information from distractors. On the other hand, if the stimulus requires a high perceptual capacity to be processed, there will not be any room for distracting information, preventing them from affecting the individual’s performance (Lavie et al., 2014).

One topic of interest within load theory literature has been how distraction is affected by the significance of the distractor. A study conducted by Biggs and colleagues (2012) has shown that search is more efficient when meaningful distractors are present, compared to meaningless ones. This conclusion is supported by studies showing that search is faster among meaningful/familiar non-targets than meaningless/unfamiliar ones (Malinowski & Hübner, 2001; Shen & Reingold, 2001). Interestingly, this does not seem to be the case for faces. It can be said that faces, due to their biological and social importance, have an additional layer of meaning for people. It was found that while pictures of fruits and instruments did not affect task performance under high perceptual load, faces were processed regardless of the perceptual load of the task (Lavie et al., 2003). This finding was explained by the fact that faces are socially important stimuli (Lavie et al., 2003).

Besides faces, the body movements of others are another important source of social information. Understanding the link between the perception of body movements and attention is crucial for human life as attention is an indispensable part of everyday social interactions. Individuals must pay attention to others in order to process and understand what they say, what they do, what they aim for, and what they mean. Studies show that this interpretation of meaning from others’ actions (i.e., action understanding) is attentionally demanding (Parasuraman & Thompson, 2012). When applied to the perceptual load theory of attention, this information means that perceiving biological motion, which is motion executed by a biological being such as a human, demands attentional sources and in turn loads the perceptual capacity of an individual. This idea is supported by previous studies that show that participants fail to perceive biological motion if their attention is captured by another stimulus (Thornton & Rensink, 2002). Therefore, it can be argued that biological motion processing requires both top-down and bottom-up mechanisms (Kroustallis, 2004). The aim of the present study is to understand how bottom-up perception of biological motion is modulated by the attentional load.

One common stimulus type utilized in biological motion studies is point-light displays (PLDs). Point-light displays (PLDs) are virtual displays of biological movements obtained from sensors placed onto joints and body parts of an individual while they are executing a certain action (Johannson, 1973). These stimuli help researchers to control many factors that could potentially influence the variables under investigation including attention. An important finding of previous biological motion studies is that the full body of the individual needs to be shown in order to clearly understand the motion (Hirai & Senju, 2020). Considering this finding, it can be concluded that understanding and processing the form of the display plays a crucial role in biological motion perception. As a result, displays that do not create a meaningful form are widely used as control conditions while studying biological motion. These displays, called scrambled motions, are created by scrambling the locations of the dots while preserving their individual motion trajectories. This way, the display is deprived of global motion and form information that creates the perception of action, while the low-level motion information of dots is maintained. Many previous behavioral and brain imaging studies have supported the idea that although the low-level characteristics are the same, biological motion and scrambled motion are processed differently (Grossman & Blake, 2001; Saygin et al., 2004). Several fMRI studies show that both ventral and dorsal pathways, which process form and motion information respectively, and superior temporal regions, where they converge, are more active when processing biological motion than scrambled motion or any other stimuli (Vaina et al., 2001; Saygin et al., 2004).

While biological motion perception has been an intense area of research in the last few decades, the majority of studies investigated it under selective attention tasks. In other words, participants usually engage in a direction discrimination or action identification task while they attend to the biological motion stimuli. Studies that investigate the bottom-up processing of biological motion stimuli while attention is directed away from them are sparse (Saygin & Sereno, 2008; Nizamoglu & Urgen, 2023). Importantly, the factors that affect the bottom-up perception of biological motion are largely unknown. The current study aims to fill these gaps by investigating the processing of task-irrelevant biological motion under varying degrees of perceptual load in three different experiments. The first experiment serves as a foundation for the following ones as it aims to study whether bottom-up processing of biological motion is affected by perceptual load and whether this processing is different from scrambled motion, which served as a control stimulus. To this end, the current study presented biological motion and scrambled motion PLDs as task-irrelevant distractors in the periphery while the perceptual load is manipulated using a letter search task in the fovea, similar to Lavie and Cox (1997). This way, it was possible to look at the extent to which biological motion is perceived under low and high perceptual load and compare it to the perception of scrambled motion. Based on the previous studies using load theory, a difference between distractor types was expected to appear in the low load condition, claiming the idea that task-irrelevant biological and non-biological motions are processed differently.

The second research question arises from the fact that each stimulus has several characteristics that influence how much attention it captures, such as its size, color, and eccentricity (Hu et al. 2020). The latter, eccentricity, refers to the distance of a stimulus from the center of the visual field. It is shown that as the eccentricity of a stimulus increases, processing speed and visual short-term memory capacity related to that stimulus tend to decrease, whereas the perceptual threshold for that stimulus increases (Staugaard et al., 2016). Additionally, there exist previous studies that extended the scope of eccentricity research by investigating the effect of eccentricity on biological motion perception. These studies show that biological motion can be perceived, and actions can be recognized up to 30° of eccentricity, after which point, recognition starts dropping to the chance level (Fademrecht et al., 2016). Moreover, the hindering effect of noise increases and sensitivity to biological motion starts to decrease as eccentricity increases (Thompson et al., 2007; Gurnsey et al., 2010). These results suggest that if biological motion is observed in the periphery, it is still possible but more difficult to process and understand the action. This effect of eccentricity on biological motion persists even when a task is present and biological motion acts as a task-irrelevant distractor (Thornton & Vuong, 2004). Although these studies investigated how biological motion perception is modulated by eccentricity, it remains unknown how it interacts with the perceptual load of a task in the fovea. Considering the previous literature on the effect of perceptual load and the effect of eccentricity, the processing of biological motion is expected to require more resources as the eccentricity of biological motion increases and the perceptual load of the task increases.

A third research question follows the second experiment. In the human brain, stimuli at different eccentricities are not processed in the same way. Cortical magnification refers to the fact that the number of neurons responsible for processing a stimulus at the center of the visual field is drastically higher than those responsible for the periphery (Duncan & Boynton, 2003). As a visual stimulus moves towards the periphery the image of the stimulus on the cortex gets proportionally smaller and consequently, a smaller number of neurons contribute to its processing. Previous studies showed that cortical magnification can modulate the effect of eccentricity on visual tasks. According to the study of Carrasco and Frieder (1997), the effects caused by eccentricity are no longer observed when the stimuli are scaled according to the cortical magnification. Johnson and Gurnsey (2010) also supported these results by showing that the sensitivity loss caused by increased eccentricity can be controlled by scaling the stimulus size. Moreover, the difference between the processing speed of peripheral and foveal stimulus decreases significantly when the peripheral stimulus is cortically scaled (Carrasco et al., 2003). However, the choice of stimuli is highly important in cortical magnification studies. For instance, the effect of eccentricity can still be strongly observed after cortical magnification if letters are used as stimuli (Staugaard, 2016). Similarly, another study has shown that the effect of eccentricity can still be seen in cortically scaled faces (Mäkelä et al., 2001). In relation to the study at hand, Gurnsey and colleagues (2008) have shown that the identification of biological motion is modulated by both its size and its eccentricity. The performance difference between the perception of foveal and peripheral biological motion can be eliminated if the stimuli are cortically scaled. (Gurnsey et al., 2008). These findings argue that the effect of cortical magnification on processing strongly depends on the task and on the stimuli. The third and final experiment aims to explore previously unanswered questions within the literature on cortical magnification. While the utilization of biological motion in such research is relatively infrequent, this experiment will investigate the interaction between perceptual load and cortical magnification on biological motion processing. Additionally, it is important to note that the biological motion stimuli will be task-irrelevant, distinguishing it from previous studies. In line with the previous studies, it is expected that when the biological motion is cortically magnified, the effects of eccentricity will be eliminated such that the stimuli at higher eccentricities will no longer require more resources.

## 2. Methods

The current study consists of three behavioral experiments. The first experiment aimed to investigate whether perceptual load affects bottom-up perception of task-irrelevant biological motion and how that effect differs from a task-irrelevant non-biological motion. The second experiment aimed to investigate if eccentricity plays a modulating role in the bottom-up perception of biological motion under different perceptual loads. The third experiment aimed to investigate if the results of the second experiment hold when cortical magnification is taken into account.

### 2.1. Experiment 1

#### 2.1.1. Participants

28 university students (19 females, mean age = 20.75; SD = 1.97) participated in the study. Participants had a normal or corrected-to-normal vision and had no history of neurological disorders. Bilkent University’s Human Research Ethics Committee approved the study and informed consent was obtained from the participants accordingly. After completing the experiment, participants were given money or course credit as compensation for their time.

#### 2.1.2. Stimuli

The experiment included two types of stimuli: task stimuli and peripheral distractor stimuli. The task stimuli consisted of six letters and are described in the Design and Procedure section. The distractor stimulus was either a PLD of a walking human or the scrambled version of the display. The PLD of the walking human (i.e. the biological motion) was selected from the CMU Graphics Lab Motion Capture Database (Carnegie Mellon University [CMU] - Graphics Lab, http://mocap.cs.cmu.edu/). By using the application “Motion Kinematic and Kinetic Analyser” ([Mokka], 2018, https://biomechanical-toolkit.github.io/mokka/), dots were removed from the stimuli until there were 13 left. These 13 dots represented the head and both sides of, shoulders, elbows, hands, waistline, legs, and feet (See Figure 1A). Then, Mokka was used once again to cut the animation into a display of 60 frames that lasted 1 second. Scrambled motion was created using a built-in function from the Biomotion Toolbox (van Boxtel & Lu, 2013), which scrambles the starting points of the dots of the walker while preserving the individual motion trajectory of each dot (See Figure 1B). This way, the global information of the human walker was eliminated, but the low-level motion properties were the same as the intact biological motion. The scrambled motion display was prepared prior to the experiment and was used for all scrambled motion trials of all subjects, in order to match the single exemplar biological motion stimulus.

**Figure 1.**
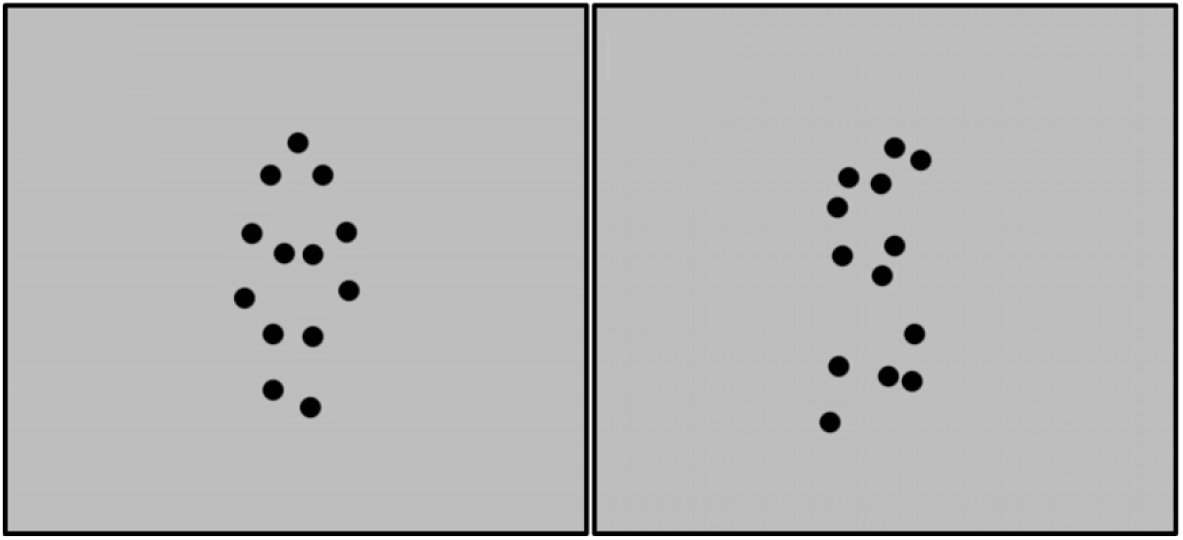
Single frames from the point-light displays consisting 13 dots. Left: Biological motion. Right: Scrambled motion.

#### 2.1.3 Design and Procedure

Upon entering the experiment room, participants were asked to sit down in front of the 21-inch monitor and place their heads on the chinrest which was used in order to ensure a fixed view distance of 56 cm in all of the trials. Before the experiment, participants were presented with biological and scrambled motion displays at the center of the screen, to familiarize them with the distractors. After that, they did a practice session of 16 trials, where they focused on the fixation at the center, and either a biological or scrambled motion was presented on the left or the right, 6 degrees away from the fixation. Participants were instructed to indicate the location of the display by pressing the left or right arrow only if the biological motion was presented. This was done to ensure that participants could differentiate between biological and scrambled motion in the periphery. Participants who performed above 85% accuracy continued to the main experiment.

The experiment and the practice session described above were programmed in MATLAB using the Psychtoolbox (Brainard, 1997; Pelli, 1997). To show the biological motion, BioMotion Toolbox was used (van Boxtel & Lu, 2013). The experiment consisted of two tasks, an easy (low perceptual load) and a hard one (high perceptual load). All participants completed both tasks in two consecutive sessions and the order of the tasks was counterbalanced across participants. Both tasks included a representation of six letters as a circle with a radius of 2.7 degrees. The task was to find and report the target letter while fixating on a fixation point that was presented at the center throughout the experiment. The target letter was always present and was either X or N. Participants were asked to press the X key on the keyboard if the present target letter was X, and N key if the present target letter was N. In the easy task, all of the non-target letters were O (See Figure 2A); whereas in the hard task, the non-target letters were similar-looking angular letters (i.e. H, W, V, K, M or Z; see Figure 2B). The relatively higher similarity of the non-target letters to the target letter during the hard task increased the perceptual load of the task, turning it into a harder one. During the experiment, each task consisted of 384 trials, half of which did not include a distractor. Of the remaining half, there were equal numbers of trials with biological and scrambled motion distractors. Each distractor appeared on the left for half of the trials, and on the right for the other half. The target letter and the target letter’s location were equally distributed across conditions, in a pseudorandom order. Before the experiment, there was a 5-trial practice session, using the task (low load or high load) the participant was going to encounter first, to make sure that they understood what they were asked to do. Before starting the experiment, the participants were instructed to respond as fast and as accurately as possible. While the central letter search task was being performed, the distractor stimulus was presented in half of the trials, at an eccentricity of 6° (see Figure 2). The letter search task, as well as the moving distractor, remained on the screen for a duration of one second. Participants were instructed to respond during this one second when the stimuli were present. During each trial, the reaction time and accuracy of the participants were recorded. There was one second interstimulus interval after each trial. Following every 128 trials, participants were allowed to take a short break. After the participants were done with the first task, they were allowed to take a longer break before the experimenter started the second task. The participants were instructed to start the task after the experimenter turned off the lights and left the room.

**Figure 2.**
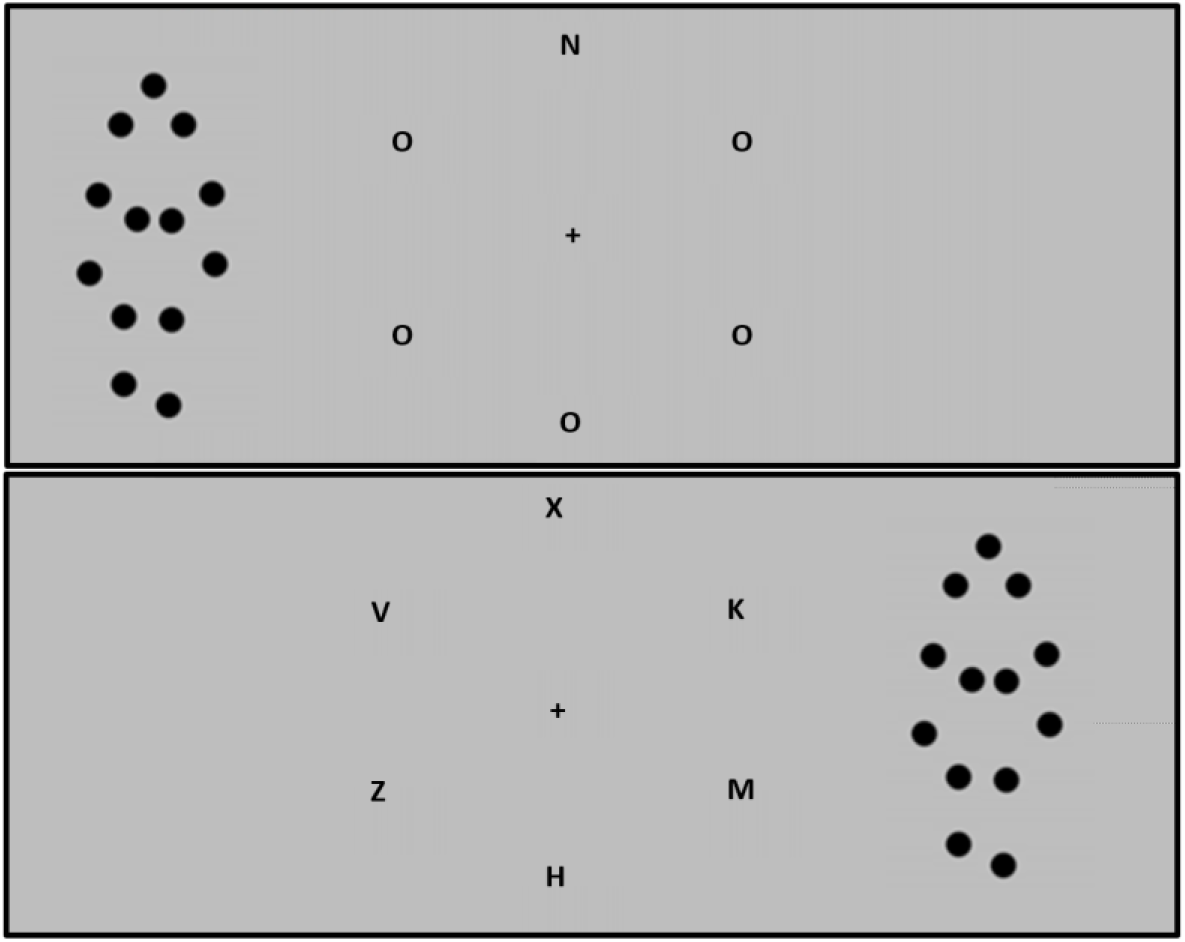
Example task display with biological motion as distractor. Top: Easy task with “N” as the target letter. Bottom: Hard task with “X” as the target letter.

#### 2.1.4. Data Analysis

Two 2x3 repeated measures ANOVAs were conducted to investigate the effect of perceptual load (low and high) and the effect of the peripheral distractor (absent, biological motion, and scrambled motion) on reaction times and accuracy.

### 2.2. Experiment 2

#### 2.2.1. Participants

31 university students (18 females, mean age = 20.71; SD = 1.59) participated in the study. Participants had a normal or corrected-to-normal vision and had no history of neurological disorders. Bilkent University’s Human Research Ethics Committee approved the study, and the informed consent was obtained from the participants accordingly. After completing the experiment, participants were given money or course credit as compensation for their time.

#### 2.2.2. Stimuli

Similar to the first experiment, Experiment 2 included two types of stimuli: (1) The task stimuli consisting of a chamber of six letters and (2) the distractor stimulus consisting of the point light display of the biological motion. They were the same as the first experiment, except that only the biological motion display was used as a distractor stimulus.

#### 2.2.3. Design and Procedure

In the second experiment, although the task of the participants was exactly the same as that of the first experiment, there were two important changes. First of all, since the participants did not have to differentiate between biological and scrambled motion, the practice session of the first experiment was not used in the second experiment. Nevertheless, participants were presented with the biological motion display before the experiment to familiarize them with the distractor. Secondly, the distractors were presented with different eccentricities. During 288 trials, the distractor was present at either 6°, 9°, or 12° away from the center, leading to 96 trials per eccentricity condition (see Figure 3). The location (left or right) of the distractor, the target letter, and the location of the target letter were equally distributed between trials in a pseudorandom order. There were also 288 trials that did not include a distractor. In total, the second experiment consisted of 576 trials. The participants were allowed to take a short break after every 144 trials, and a longer break between tasks, similar to the first experiment.

**Figure 3.**
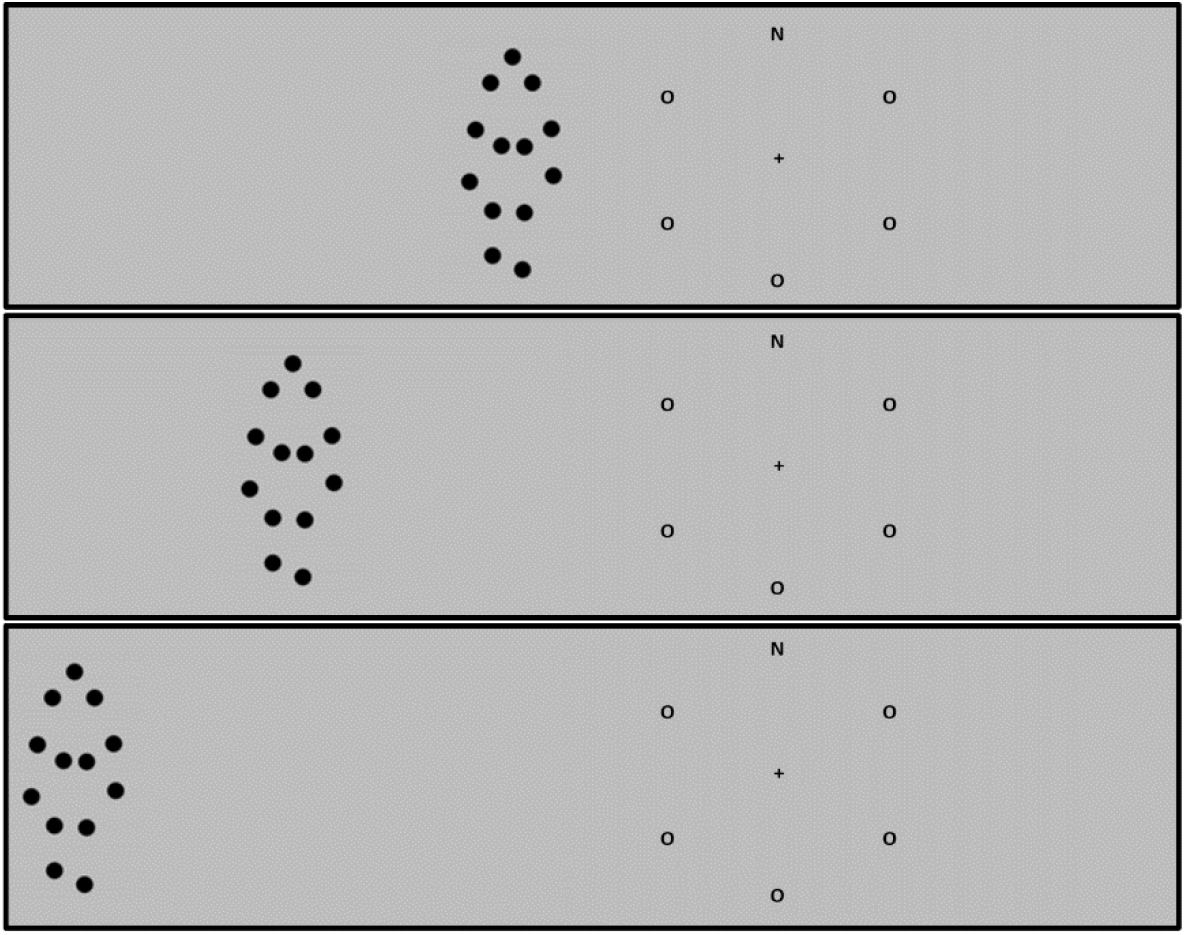
Example trials from Experiment 2. Top: Example trial for 6° condition. Middle: Example trial for 9° condition. Bottom: Example trial for 12° condition.

#### 2.2.4. Data Analysis

Two 2x4 repeated measures ANOVAs were conducted to investigate the effect of perceptual load (low and high) and the effect of the peripheral distractor (with the levels of absent, 6 degrees, 9 degrees, and 12 degrees) on reaction times and accuracy.

### 2.3. Experiment 3

#### 2.3.1. Participants

36 university students (25 females, mean age = 21.72; SD = 2.06) participated in the study. Participants had a normal or corrected-to-normal vision and had no history of neurological disorders. Bilkent University’s Human Research Ethics Committee approved the study, and informed consent was obtained from the participants accordingly. After completing the experiment, participants were given money or course credit as compensation for their time.

#### 2.3.2 Stimuli

The task and the distractor stimuli were the same as in the second experiment. However, the distractor stimulus (the biological motion display) was scaled in size proportionally across different eccentricity conditions. The stimulus size was the same as that in Experiment 2 for the 6° condition, whereas its size was scaled by x1.5 and by x2 for 9° and 12° conditions, respectively.

#### 2.3.3 Design and Procedure

The design and procedure were exactly the same as the second experiment. The scaled biological motions were used as the distractor stimuli (see Figure 4).

**Figure 4.**
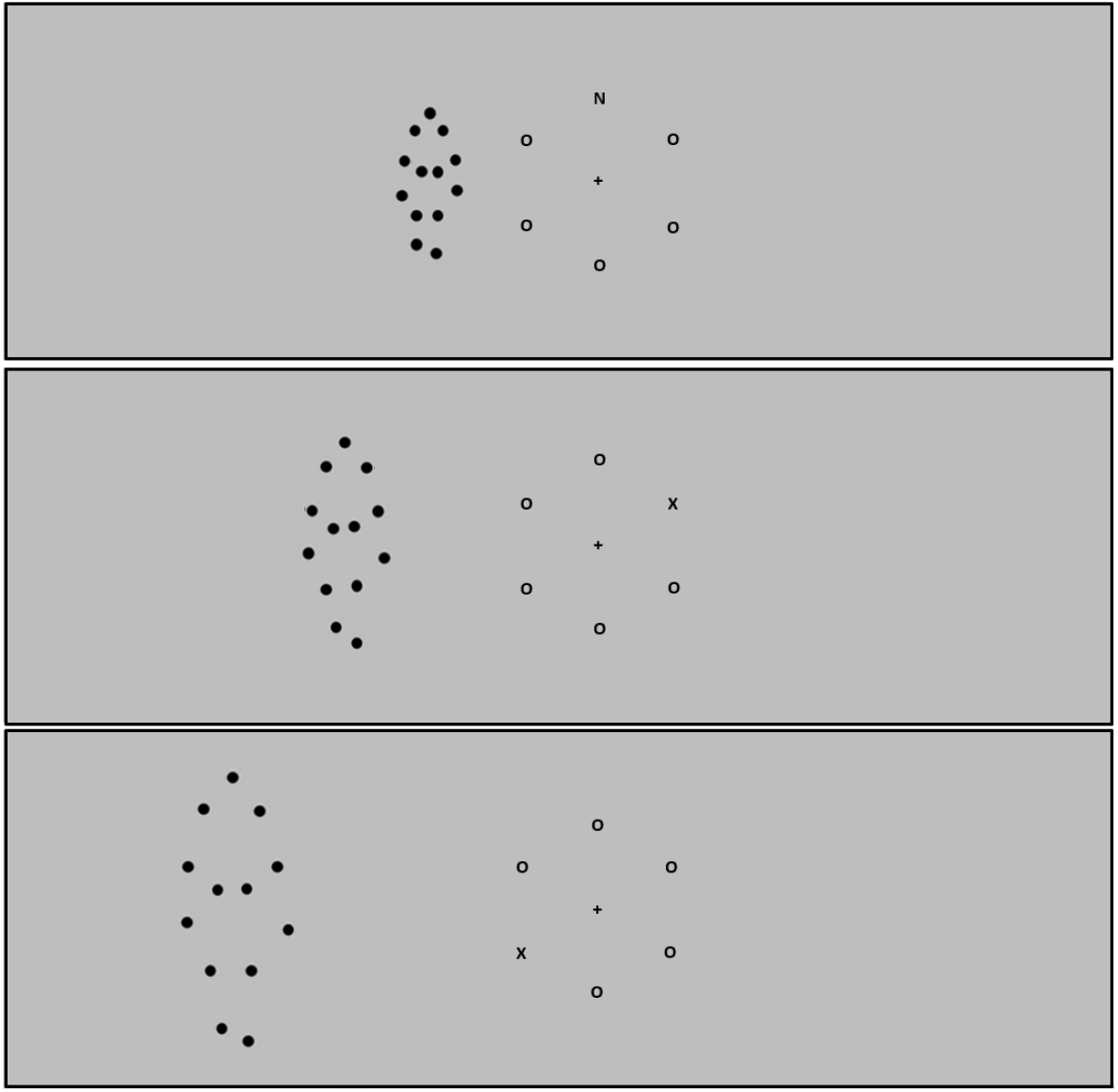
Example trials from Experiment 3 where the peripheral stimulus size is scaled proportionally with the eccentricity. Top: Example trial for 6° condition. Middle: Example trial for 9° condition. Bottom: Example trial for 12° condition.

#### 2.3.4 Data Analysis

Two 2x4 repeated measures ANOVAs were conducted to investigate the effect of perceptual load (low and high) and the effect of the distractor (absent, 6 degrees, 9 degrees and 12 degrees) on the reaction times and on the accuracy.

## 3. Results

The missed trials from all three of the experiments are removed from the data for the purposes of reaction time and accuracy analyses.

### 3.1 Experiment 1

#### 3.1.1 Reaction Time

Reaction time analysis revealed main effects of perceptual load (F(1,27) = 432.320, p<0.001, η²_p_ = 0.941) and distractor type (F(2,54) = 238.777, p<0.001, η²_p_ = 0.898). Participants were significantly slower in the hard task with high perceptual load (M = 0.738, SE = 0.011) compared to the easy task with low perceptual load (M = 0.546, SE = 0.011). Additionally, participants were significantly slower in the scrambled motion condition (M = 0.658, SE = 0.010) compared to the biological motion condition (M = 0.651, SE = 0.010) (t(27) = 3.199, p<0.05). In the easy task, participants were faster while there was no distractor (M = 0.521, SE = 0.011), compared to biological motion (M = 0.555, SE = 0.012) (t(27) = 12.362, p<0.001) and scrambled motion (M = 0.561, SD = 0.012)(t(27) = 14.326, p<0.001) conditions. Moreover, the results have shown that the participants reacted faster in the biological motion condition compared to the scrambled motion condition. This effect was approaching significance (t(27) = 1.963, p = 0.052). In the hard task, participants were faster when there was no distractor (M = 0.711, SE = 0.011) compared to when there was a distractor of biological motion (M = 0.747, SE = 0.011)(t(27) = 13.122, p<0.001) and scrambled motion (M = 0.755, SE = 0.012)(t(27) = 15.920, p<0.001). Additionally, participants reacted faster in the biological motion condition compared to the scrambled motion condition (t(27) = 2.798, p = 0.012). Lastly, a significant interaction between perceptual load and distractor type was not observed (F(2,54) = 0.713, p>0.05).

**Figure 5.**
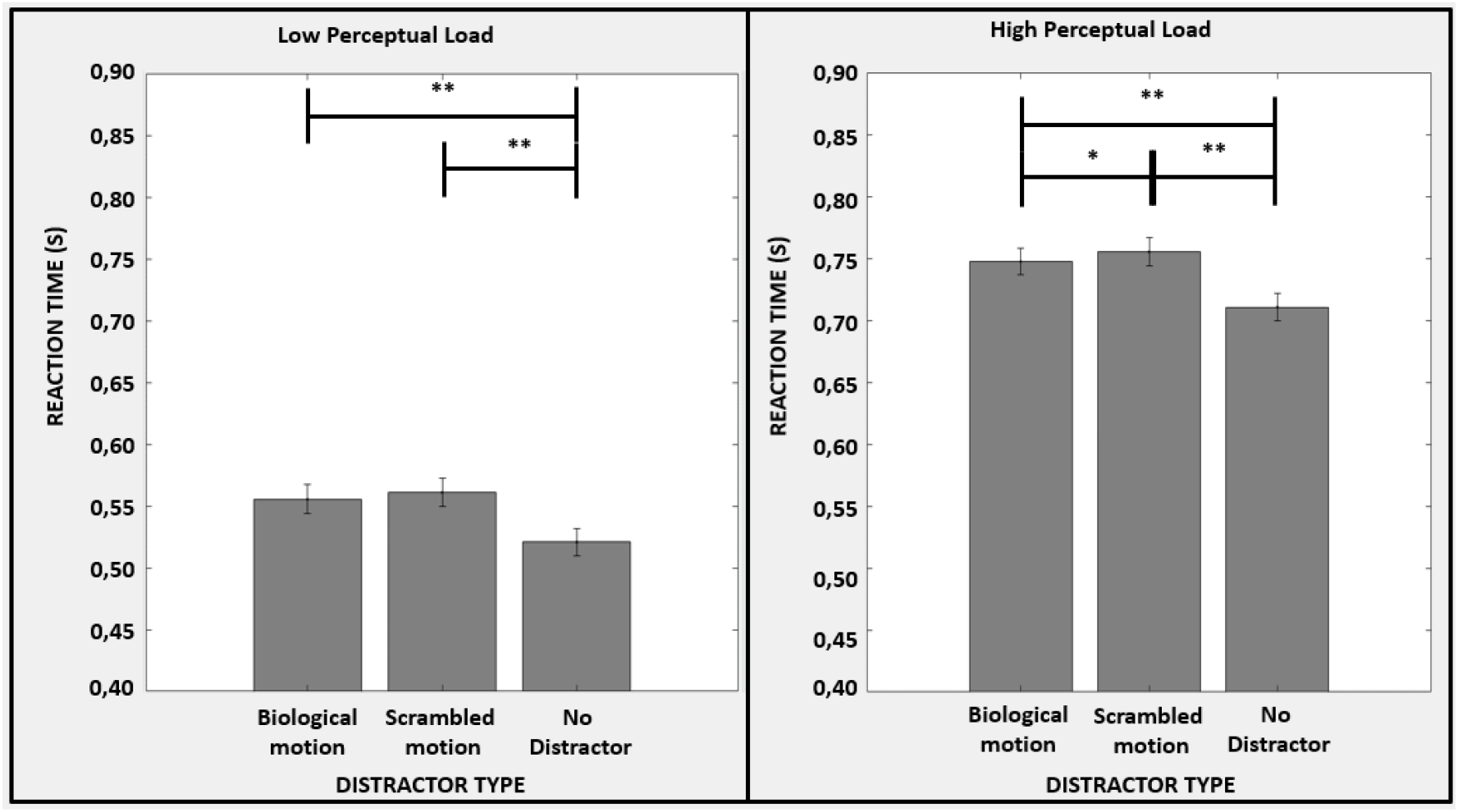
Reaction time data that represents low load (top) and high load (bottom) conditions for the second experiment (*: p<0.05, **: p<0.001).

#### 3.1.2 Accuracy

Accuracy analysis revealed a main effect of perceptual load (F(1,27) = 81.022, p<0.001, η²_p_ = 0.750) whereas the effect of distractor type was not significant (p>0.05). Participants were significantly more accurate in the easy task (M = 95.4%, SE = 1.3%) compared to the hard task (M = 82.4%, SE = 1.3%) (t(26) = 9.001, p<0.001). In the easy task, participants were equally accurate in no distractor (M = 95.2%, SE = 0.6%), biological motion (M = 95.1%, SE = 0.7%), and scrambled motion (M = 95.9%, SE = 0.6%) conditions (p>0.05 for all). Similarly, in the hard task, participants were equally accurate in no distractor (M = 83.0%, SE = 1.7%), biological motion (M = 82.2%, SE = 1.9%), and scrambled motion (M = 82.0%, SE = 1.9%) conditions (p>0.05 for all). Lastly, there was no significant interaction effect between the distractor and the perceptual load (p>0.05).

**Figure 6.**
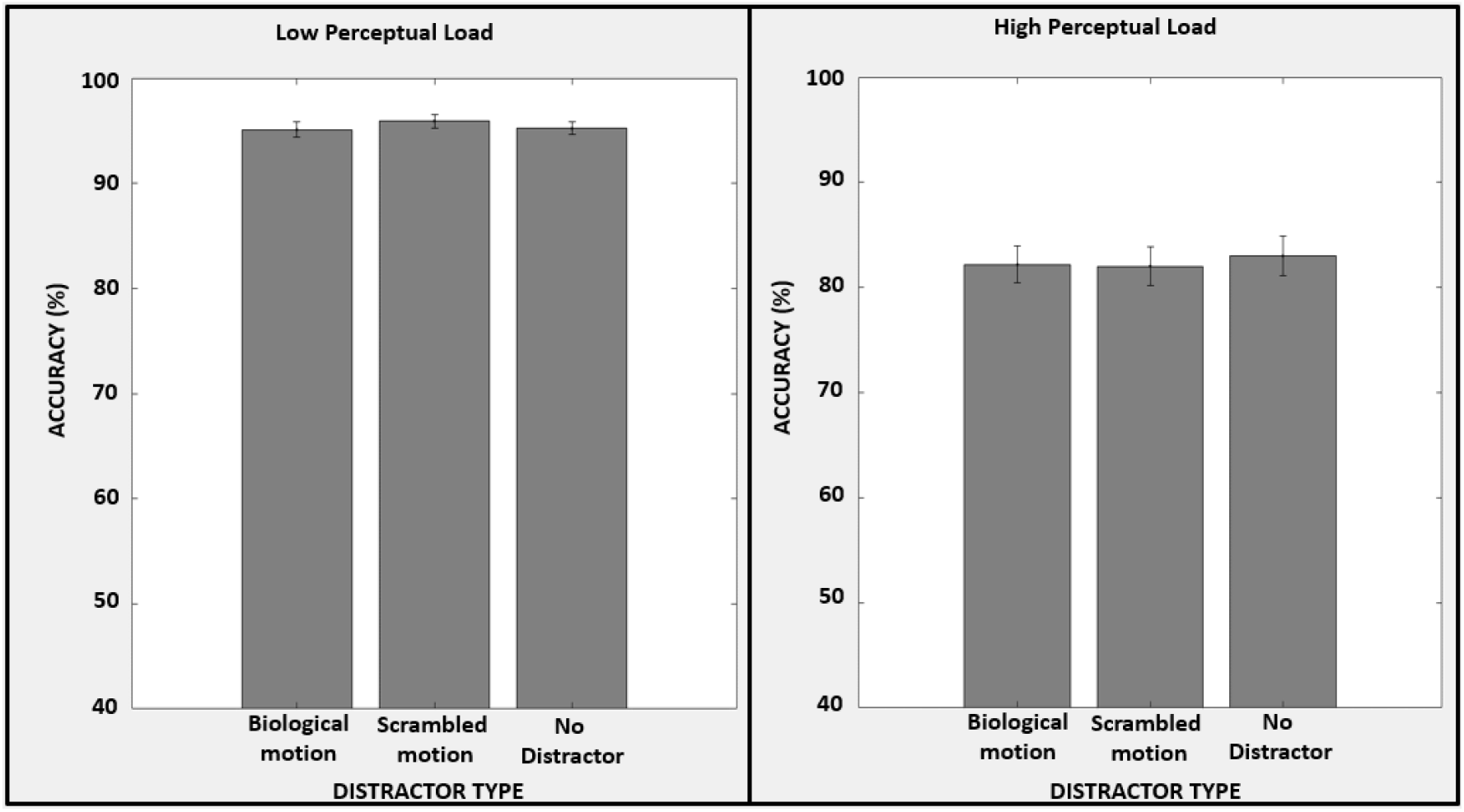
Accuracy data that represents low load (top) and high load (bottom) conditions for the first experiment.

### 3.2 Experiment 2

Four outliers have been detected and excluded from further analysis according to the interquartile range rule. Three of them performed poorly in the low-load task with accuracy rates of 88%, 85%, and 82%. The last outlier was excluded from the analysis as they missed 87.5% of the trials in the high-load task.

#### 3.2.1 Reaction Time

Reaction time analysis revealed main effects of perceptual load (F(1,26) = 482.526, p<0.001, η²_p_ = 0.914) and distractor type (F(3,78) = 98.109 p<0.001, η²_p_ = 0.791). Participants were significantly slower in the hard task with high perceptual load (M = 0.756, SE = 0.008) compared to the easy task with low perceptual load (M = 0.563, SE = 0.008)(t(26) = 21.966, p<0.001). In the easy task, participants were faster while there was no distractor (M = 0.536, SE = 0.009), compared to the 6° (M = 0.579, SE = 0.009)(t(26) = 11.911, p<0.001), 9° (M = 0.568, SE = 0.009)(t(26) = 8.752, p<0.001) and 12° (M = 0.568, SE = 0.009)(t(26) = 8.731, p<0.001) conditions. Moreover, the 6° condition was significantly slower than 9° (t(26) = 3.160, p<0.05) and 12° (t(26) = 3.180, p<0.05) conditions. No difference was observed among 9° and 12° conditions (p>0.05). In the hard task, participants were faster when there was no distractor (M = 0.729, SE = 0.007) compared to the 6° (M = 0.765, SE = 0.009)(t(26) = 10.007, p<0.001), 9° (M = 0.764 SE = 0.007)(t(26) = 9.712, p<0.001) and 12° (M = 0.765, SE = 0.007)(t(26) = 9.967, p<0.001) conditions. Additionally, no difference was observed among 6°, 9°, and 12° conditions (p>0.05 for all). Lastly, there was no significant interaction effect between the distractor and the perceptual load (p>0.05).

**Figure 7.**
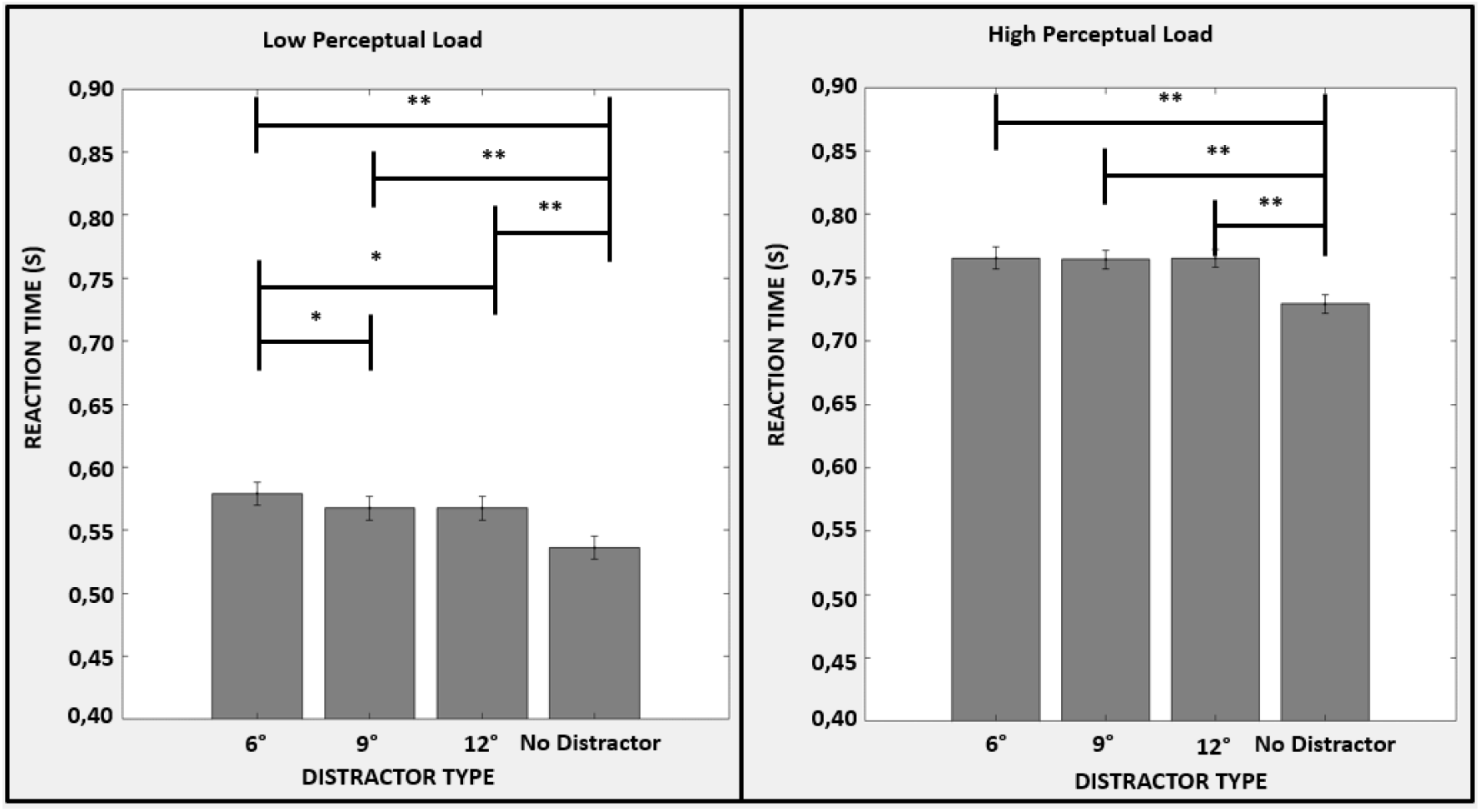
Reaction time data that represents low load (top) and high load (bottom) conditions for the second experiment (*: p<0.05, **: p<0.001).

#### 3.2.2 Accuracy

Accuracy analysis revealed a main effect of perceptual load (F(1,26) = 128.213, p<0.001, η²_p_ = 0.831) whereas the effect of distractor type was not significant (p>0.05). Participants were significantly more accurate in the easy task (M = 96.7%, SE = 0.9%) compared to the hard task (M = 85.2%, SE = 0.9%)(t(26) = 11.323, p<0.001). In the easy task, participants were equally accurate in no distractor (M = 96.9%, SE = 0.4%), 6° (M = 96.7%, SE = 0.5%), 9° (M = 96.3%, SE = 0.6%) and 12° (M = 96.9% SE = 0.5%) conditions (p>0.05 for all). Similarly, in the hard task, participants were equally accurate in no distractor (M = 86.3%, SE = 1.1%), 6° (M = 84.0%, SE = 1.5%), 9° (M = 84.9%, SE = 1.5%) and 12° (M = 85.5% SE = 1.3%) conditions (p>0.05 for all). Lastly, there was no significant interaction effect between the distractor and the perceptual load (p>0.05).

**Figure 8.**
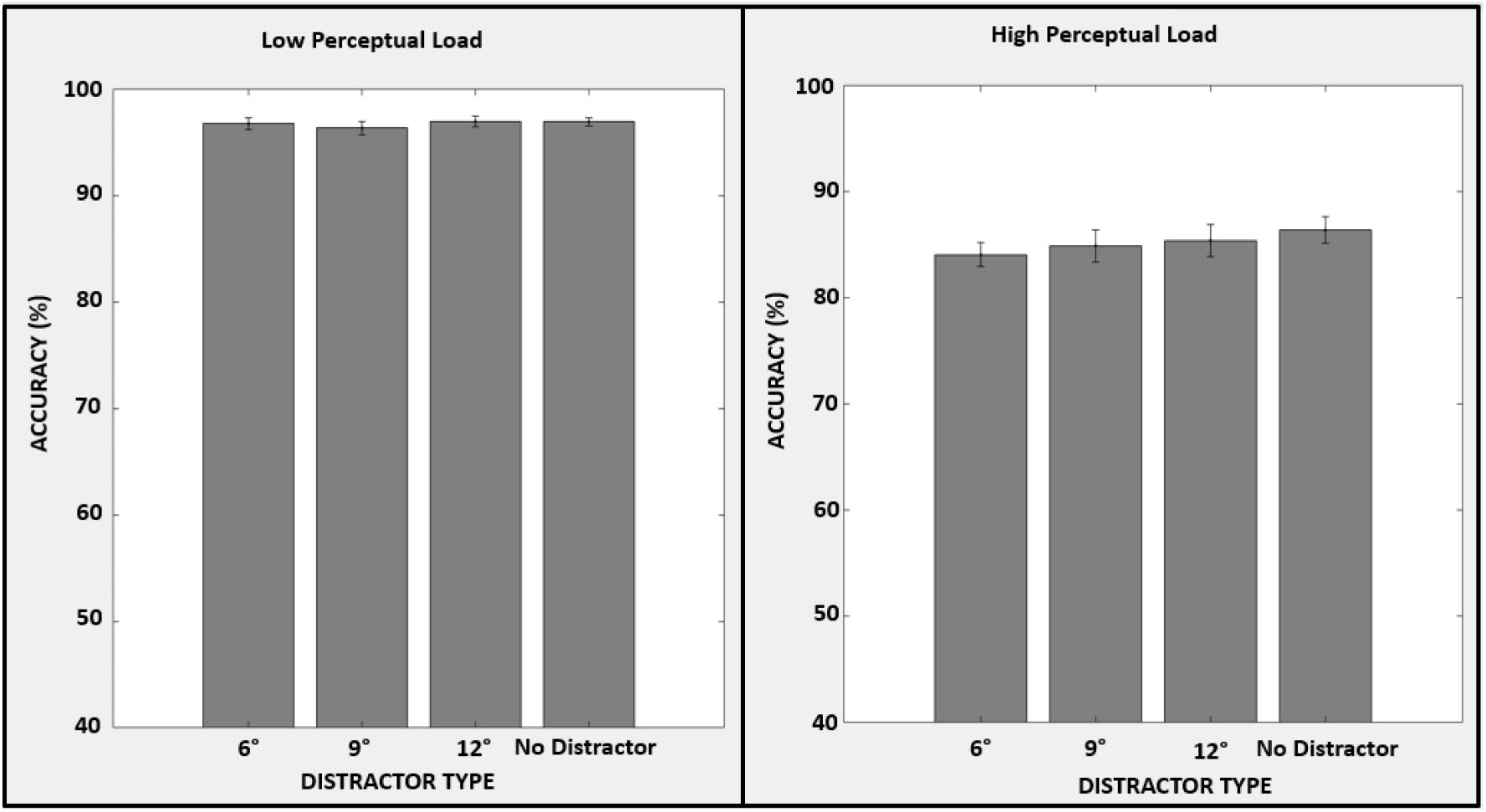
Accuracy data that represents low load (top) and high load (bottom) conditions for the second experiment.

### 3.3 Experiment 3

Five outliers have been detected and excluded from further analysis according to the interquartile range rule. Two of them performed poorly in the low load task with accuracy rates of 85% and 88%, one of them performed poorly in the high load task with an accuracy rate of 46%, and the remaining two missed 49.5% and 54.5% of the trials in the high load task.

#### 3.3.1 Reaction Time

Reaction time analysis revealed main effects of perceptual load (F(1,30) = 403.136, p<0.001, η²_p_ = 0.931) and distractor type (F(3,90) = 113.038 p<0.001, η²_p_ = 0.790). Participants were significantly slower in the hard task with high perceptual load (M = 0.729, SE = 0.008) compared to the easy task with low perceptual load (M = 0.538, SE = 0.008)(t(30) = 20.078, p<0.001). In the easy task, participants were faster while there was no distractor (M = 0.514, SE = 0.007), compared to the 6° (M = 0.552, SE = 0.008)(t(30) = 12.808, p<0.001), 9° (M = 0.543, SE = 0.007)(t(30) = 9.725, p<0.001) and 12° (M = 0.541, SE = 0.007)(t(30) = 9.128, p<0.001) conditions. Moreover, the 6° condition was significantly slower than 9° (t(30) = 3.083, p<0.05) and 12° (t(30) = 3.680, p<0.01) conditions. No difference was observed among 9° and 12° conditions (p>0.05). In the hard task, participants were faster when there was no distractor (M = 0.723, SE = 0.008) compared to the 6° (M = 0.749, SE = 0.009)(t(30) = 8.753, p<0.001), 9° (M = 0.757 SE = 0.008)(t(30) = 11.465, p<0.001) and 12° (M = 0.751, SE = 0.009)(t(30) = 9.423, p<0.001) conditions. Additionally, 6° condition yielded significantly faster reaction times compared to 9° condition (t(30) = 2.712, p<0.05) although no difference was observed among 6° and 12° conditions or 9° and 12° conditions (p>0.05 for all). Lastly, a significant interaction effect was observed between the distractor and the perceptual load (F(3,90) = 5.826, p<0.01, η²_p_ = 0.163).

**Figure 9.**
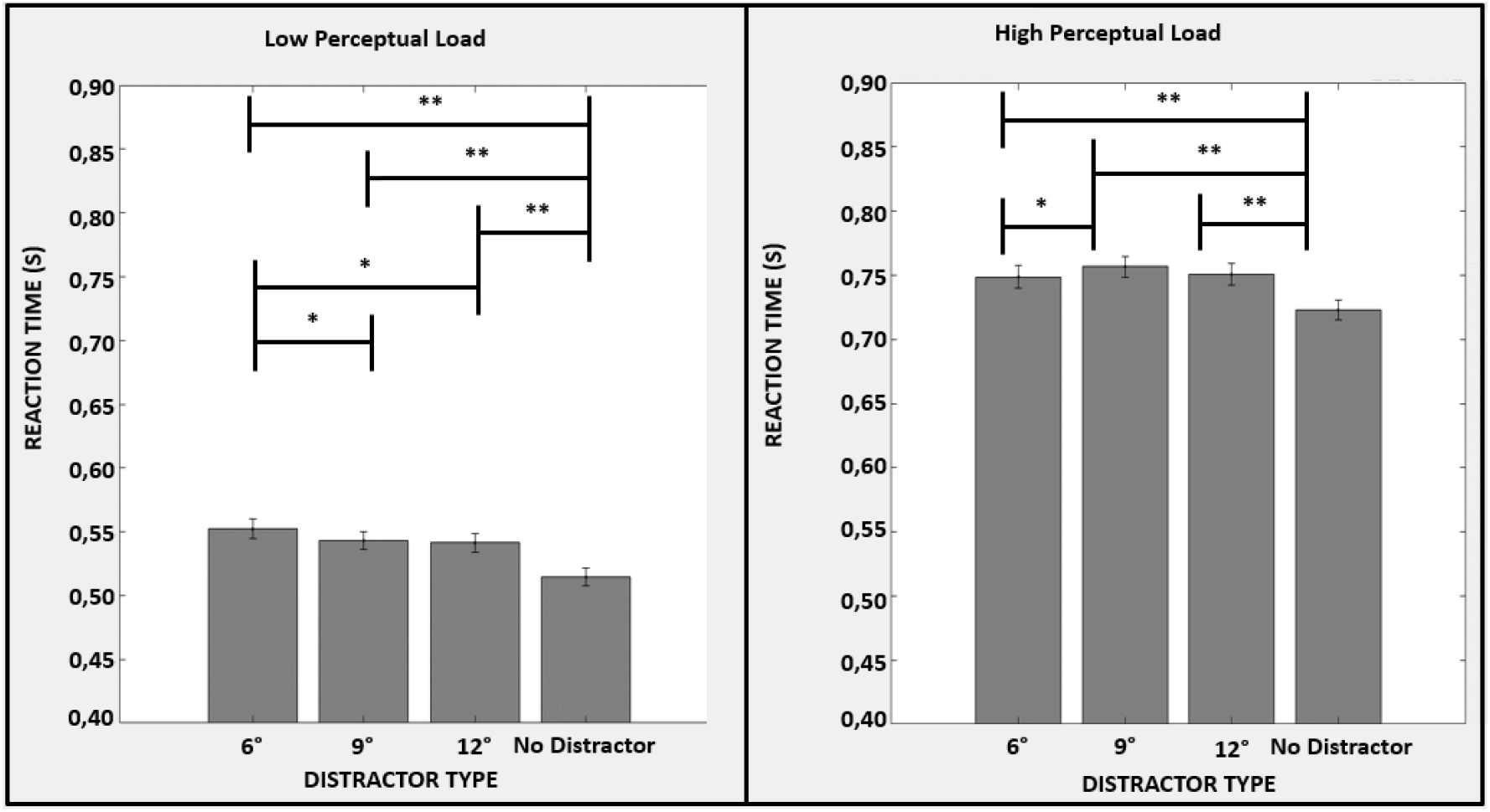
Reaction time data that represents low load (top) and high load (bottom) conditions for the third experiment (*: p<0.05, **: p<0.001).

#### 3.3.2 Accuracy

Accuracy analysis revealed a main effect of perceptual load (F(1,30) = 129.813, p<0.001, η²_p_ = 0.812) and distractor type (F(3,90) = 4.353, p<0.01, η²_p_ = 0.127). Participants were significantly more accurate in the easy task (M = 95.3%, SE = 0.4%) compared to the hard task (M = 83.9%, SE = 1.2%)(t(30) = 11.394, p<0.001). In the easy task, participants were equally accurate in no distractor (M = 95.4%, SE = 0.4%), 6° (M = 95.2%, SE = 0.6%), 9° (M = 95.4%, SE = 0.5%) and 12° (M = 95.1%, SE = 0.6%) conditions (p>0.05 for all). In the hard task, participants were significantly more accurate in the no distractor (M = 85.5% SE = 1.1%) condition compared to 9° (M = 82.0%, SE = 1.3%)(t(30) = 4.641, p<0.001) condition. Additionally, participants were significantly less accurate in 9° condition compared to 12° (M = 84.9%, SE = 1.3%)(t(30) = 3.859, p<0.05) condition. No difference was observed among 6° (M = 83.4%, SE = 1.5%) and 9° conditions or 6° and 12° conditions (p>0.05 for all). Moreover, no difference was observed among no distractor and 12° or no distractor and 6° conditions as well. (p>0.05). Lastly, a significant interaction effect was observed between the distractor and the perceptual load (F(3,90) = 4.415, p<0.01, η²_p_ = 0.128).

**Figure 10.**
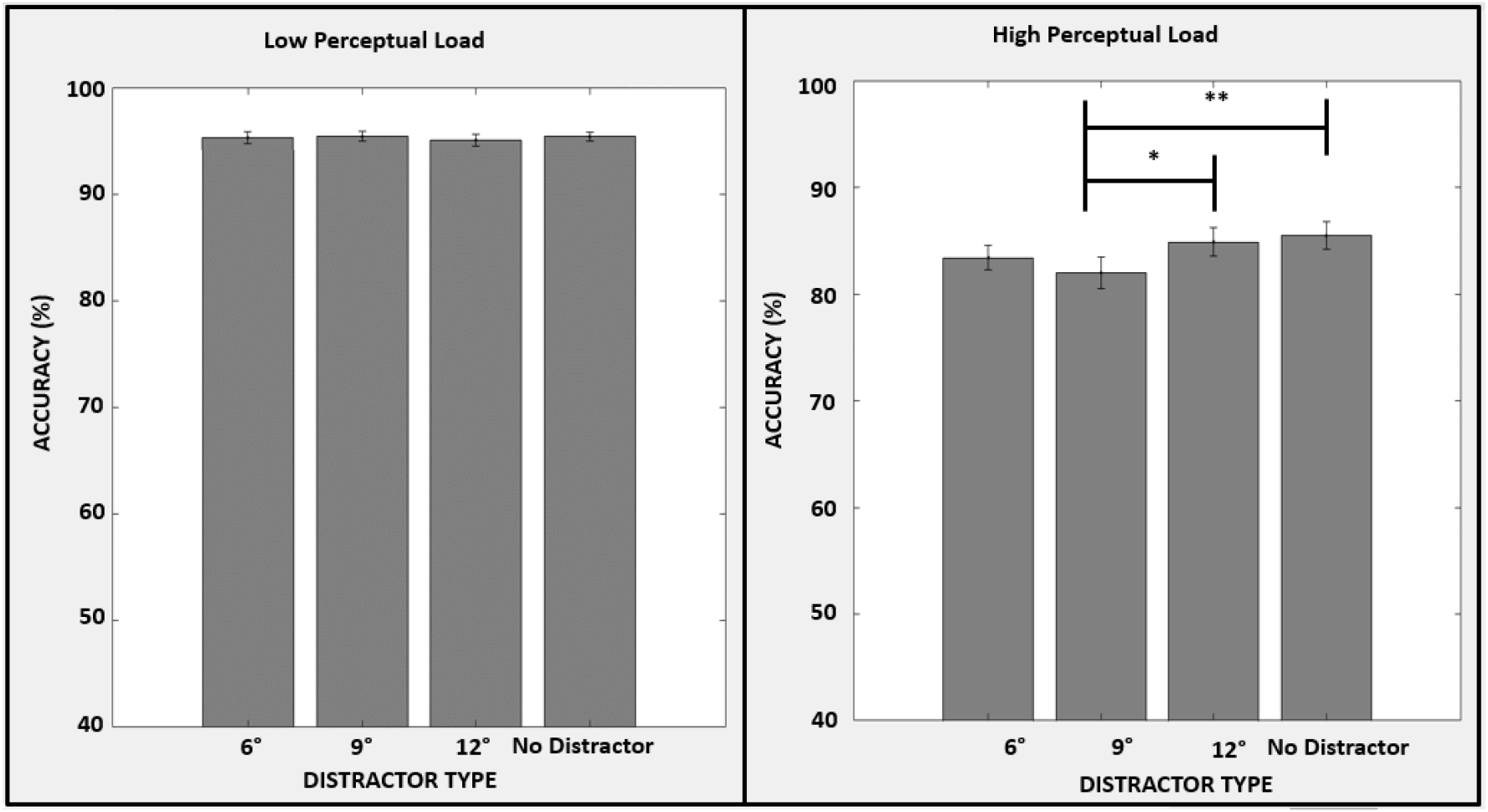
Accuracy data that represents low load (top) and high load (bottom) conditions for the third experiment (*: p<0.05, **: p<0.001).

## 4. Discussion

Over the course of three experiments, the current study investigated the influence of perceptual load on the bottom-up perception of biological motion. In the first experiment, we establish that even when presented as a task-irrelevant peripheral distractor, biological motion is processed differently than scrambled motion. More specifically, we found that the reaction times of the participants became slower when a scrambled motion was presented as the distractor, compared to the biological motion distractor. This finding suggests that the processing of scrambled motion is more attentionally demanding than biological motion. Moreover, this effect is significant when the perceptual load is high, and approaches significance when the perceptual load is low. The second experiment was then conducted to investigate the effect of eccentricity (6°, 9°, and 12°) on the bottom-up perception of biological motion under different levels of perceptual load. It expands the findings of the first one by suggesting that the processing of biological motion is modulated by its eccentricity. We found that as the eccentricity increases, the stimulus becomes less attentionally-demanding. However, this effect seems to be present only when the perceptual load is low and within a certain eccentricity range between 6° and 9°. Lastly, the third experiment investigated whether the results of the second experiment would hold when the cortical magnification factor was taken into account. The results show that if the stimuli at different eccentricities are scaled in line with the cortical magnification factor, biological motion becomes more attentionally-demanding when the perceptual load is high and the eccentricity is 9°. Additionally, all three experiments show the main effect of the perceptual load as participants performed significantly better in the low-load task than in the high-load task in terms of reaction time and accuracy.

### The effect of perceptual load on the bottom-up perception of biological motion

The first experiment supports and expands the results of previous experiments investigating the attentional load theory. As predicted by the theory, longer reaction times and lower levels of accuracy were observed for the high-load task than for the low-load task. Although it is still possible to process biological motion distractors during the high-load task, their effect on attention is considerably less because of the lower amount of capacity left compared to the easy task. Moreover, the reaction times were increased when a distractor was present, compared to the no distractor condition under both perceptual loads. This indicates that both biological and scrambled motion when presented in the periphery as task-irrelevant distractors, are capable of capturing attention in a bottom-up fashion and impair performance in a central task. These results highlight the significance of the interplay of bottom-up and top-down factors in biological motion perception and necessitate the extension of solely feedforward models of biological motion perception (Giese and Poggio, 2003) by incorporating top-down and bottom-up attention in computational modeling studies.

An important finding of the first experiment is that the scrambled motion distractor interfered with task performance more than the biological motion distractor as revealed by increased reaction times in the former compared to the latter. However, there was no significant difference in accuracy between biological and scrambled motion distractors. This can be explained by the speed-accuracy tradeoff. Participants, instead of giving quick but inaccurate responses, tend to favor being slow and accurate (Dean et al., 2007). Overall, these findings are in line with studies that suggest that search is more efficient among distractors that are meaningful or familiar, compared to those that are not. A possible explanation is that familiar and meaningful stimuli get processed and categorized faster (Mruczek & Sheinberg, 2005; Biggs et al., 2012), which would lead to faster rejection of distractors. According to this explanation, in the context of our experiment, when the distractor is presented with the letter search task display, it may initially capture attention, which is dedicated to processing and identifying it. Once it is processed, it can be categorized (e.g., as a walker) and then rejected, as it is not the object of the search. Since biological motion contains familiar global motion information, it can be categorized faster and rejected accordingly. However, scrambled motion cannot be identified and fit into a stimulus category. Thus, it may employ attentional resources for a longer time before rejection, which may take away from the resources that can be used for the task. A second possible explanation is that distractor familiarity may affect pre-attentive processes (Wolfe, 2001). This account would suggest that the novelty of a stimulus can be a pre-attentive feature that can lead to further attentive processing. Due to its unfamiliar form and motion, scrambled motion would be pre-attentively categorized as novel, which would cause attentional resources to be allocated to it, and away from the main task. Since biological motion is more familiar (less novel), it could capture fewer resources, leading to the pattern of distractor interference observed in the current study.

### The effect of eccentricity on the bottom-up perception of biological motion under perceptual load

The results of the second experiment support the first one such that the participants reacted faster and more accurately in the easy task compared to the hard task. Moreover, similar to the previous experiment, the presence of a distractor caused an increase in the reaction times compared to the no distractor condition. Importantly, the second experiment built up on the results of the first experiment by showing that the bottom-up processing of biological motion not only depends on perceptual load but also eccentricity. Our results show that as the eccentricity of the biological motion decreases, the reaction times get higher. This finding is consistent with previous work (Thornton & Vuong, 2004). However, no difference was observed in accuracy levels across different eccentricities, which can possibly be explained by the speed-accuracy tradeoff (Dean et al., 2007). On the other hand, our results also indicate an interaction between eccentricity and perceptual load. We found that distractors at lower eccentricities increase reaction times only during the low-load task. The lack of a difference in reaction times across different eccentricities during the high-load task can be explained by the fact that processing distractors at distant eccentricities away from the central visual field requires a greater amount of attentional resources compared to distractors at close eccentricities. When the perceptual load is low, there is enough perceptual capacity left to process the stimuli even at distant eccentricities. However, when the perceptual load is high, not enough perceptual capacity is left to process the stimuli at distant eccentricities. These results are consistent with some previous fMRI studies (Schwartz et al, 2004). On the other hand, it is important to acknowledge that there seems to be an eccentricity range (up to 9°) in which biological motion is distracting, and beyond this range, it is not distracting as much. This leads us to consider the possibility of an eccentricity threshold in the distractibility of the biological motion stimuli. This current experiment is unable to locate the exact value of this threshold (i.e., location away from the central visual field in visual degrees) and future studies should address this by investigating the changes in the reaction time for eccentricity values between 6 and 9 degrees.

### Taking into account the cortical magnification factor: The effect of eccentricity on the bottom-up perception of biological motion under perceptual load

The aim of the third experiment was to investigate whether cortical magnification would eliminate the differences in the perception of task-irrelevant biological motion distractors across the visual field and whether this effect depends on the perceptual load on the fovea. Together with the results of the second experiment, our results suggest that cortical magnification does not eliminate the differences in the perception of task-irrelevant biological motion distractors across the visual field when the perceptual load is low. However, when the perceptual load is high, cortical magnification brings back differences in the eccentricity that originally do not exist for the cortically non-magnified stimuli under a high perceptual load.

These results at first seem inconsistent with previous work on cortical magnification. Many behavioral and neural studies show that cortical magnification eliminates the differences in perceiving stimuli at different eccentricities (Carrasco & Frieder, 1997; Carrasco et al., 2003; Jigo et al., 2023). However, our study differs from those in several aspects. First, previous studies usually utilize paradigms in which the cortically magnified stimuli across the visual field (i.e., at different eccentricities) are task-relevant. In other words, top-down attention is usually exerted on the peripheral stimuli. On the other hand, in our study, the cortically magnified stimuli are task-irrelevant and presented as peripheral distractors. Therefore, the processing of these stimuli requires bottom-up attention. Moreover, our paradigm also takes into account the perceptual load in the fovea, so the processing of the peripheral distractors also depends on how much perceptual capacity is left from the central task. Thus, our results suggest that cortical magnification differentially affects the top-down and bottom-up processing of stimuli across the visual field and interacts with the perceptual load in the fovea.

One possible explanation for the differences in the effect of cortical magnification between the stimuli that require top-down attention and bottom-up attention is that top-down attention enhances the processing of the stimuli regardless of their position in the visual field (Sawaki & Katayama, 2008; Noudoost et al., 2010). This provides an advantage over bottom-up attention since there is a larger population of neurons that process the stimuli in the periphery for the cortically-magnified stimuli (Jigo et al., 2023) whose activities are enhanced by attention (Saygin & Sereno, 2008). Thus, our results in the context of previous work suggest that the elimination of differences in perceptual sensitivities across the visual field with cortical magnification may not solely depend on the increasing number of neurons (i.e., the spatial extent of activation on the cortex) but also on the enhancement of the activity of these neurons by top-down attention.

Furthermore, the interaction of cortical magnification with perceptual load in the fovea provides some novel insights into the visual processing of peripheral stimuli. There is mounting evidence including the results of this study that suggest that task-irrelevant peripheral stimuli may not be processed efficiently when there is not enough perceptual capacity due to a high-load task in the fovea (Pugnaghi et al., 2020). Our results extend this finding and indicate that when the peripheral stimuli are cortically magnified, their processing is enhanced even if the perceptual load in the fovea is high. This suggests that when the distracting capacity of a peripheral stimulus is strong enough (by means of cortical magnification in our case), it may cause a more flexible allocation of perceptual capacity than it is suggested by Lavie (1995)’s load theory.

### Limitations

While interpreting the results of this study, it is important to acknowledge its limitations as well. There exist several factors that might possibly affect the results, most notably the selection of stimulus. In line with the previous biological motion studies, the point-light display of a walker was the stimulus of preference for the experiments. However, the results might differ for different kinds of biological motion stimuli. For instance, presenting a display in which a certain gender, age, or mood can be identified may potentially give different results. Additionally presenting a communicative action (such as waving) may distract the participants more than walking, which is a non-communicative action. Therefore, further studies can be conducted by changing the stimulus type to increase the results’ generalizability. Another possible factor that might have affected the results is that no eye tracking was present during the experiment. Even though the participants were strictly instructed to only fixate at the fixation point, they were not monitored to make sure that they followed this instruction, except during the practice period. To that end, the study can be replicated by using an eye tracker.

### Future directions and conclusions

Besides the limitations, the study brings further questions that can be addressed in future studies. Brain imaging techniques can be used to have a neural understanding of the effect of perceptual load, eccentricity, and cortical magnification when biological motion is presented in the periphery as a task-irrelevant stimulus. Overall, the current study contributed to the literature by investigating the factors that affect the processing of a highly socially prevalent stimulus, while also opening possibilities for many further studies.

## Acknowledgments

We would like to thank our research assistants who helped us with the data collection: Ayşesu İzgi, Ece Altınbaş, Zelal Eltaş.

## Notes

### Competing Interest Statement

The authors have declared no competing interest.

